# Rats and axolotls share a common molecular signature after spinal cord injury enriched in collagen-1

**DOI:** 10.1101/184713

**Authors:** Athanasios Didangelos, Katalin Bartus, Jure Tica, Michele Puglia, Bernd Roschitzki, Elizabeth J. Bradbury

**Author notes:** Correspondence: A Didangelos.

## Abstract

Spinal cord injury (SCI) in mammals leads to irreversible tissue damage and loss of function. In contrast, axolotls are able to regenerate scar-free the injured spinal cord. To explore new pathological mechanisms, we compared rat versus axolotl transcriptomics and isolated genes shared between species post-SCI. Unexpectedly, multiple transcripts involved in extracellular matrix remodelling, in particular collagen-1, were upregulated in both species after SCI. Proteomics validated persistent expression of the collagen-enriched matrix signature at the protein level. Collagen-1 accumulated in early and advanced rat lesions. Importantly, collagen-1 was likely associated with pathological vascular remodelling rather than classic fibrosis and the transcription factor SP1 was predicted and validated to regulate, at least in part, the expression of collagen-1 in rat lesions.

## INTRODUCTION

Severe injuries to the mammalian spinal cord are characterised by lack of positive wound healing and regeneration and often result in permanent functional impairment (1). Following SCI, chronic inflammation and reactive gliosis drive neuronal loss and irreversible tissue scarring (2, 3). The maladaptive inflammatory response and accumulation of extracellular matrix (ECM) proteins in lesions, especially large chondroitin-sulfate proteoglycans (4, 5), are the major pathological determinants and are largely responsible for the lack of neuronal regeneration after SCI (6, 7). Although significant advancements have been made in understanding pathological as well as regenerative mechanisms in SCI, there are no therapies that can promote complete tissue repair.

In contrast to mammals, amphibian urodela such as axolotl mexicanum have the ability to fully regenerate most tissues including the spinal cord (8, 9). While the mechanisms behind this remarkable ability are not entirely understood, previous work points towards the combined molecular effect of progenitors, excellent patterning and a very effective immune response (10-12). Mammalian SCI has not been compared extensively to axolotls, yet such comparisons have the potential to identify conserved molecular mechanisms with unknown pathological or regenerative potential in SCI (13).

## METHODS

### Rat and Axolotl microarray data

Rat T7 spinal clip compression injury (7 days) microarray (14) was obtained from GEO-NCBI (**goo.gl/Hz4Gj9**). Axolotl mexicanum spinal cord injury microarray (9) data was obtained from the same server (**goo.gl/Eoshgn**). More details in **Supplemental Methods.**

### SCI model

Isofluorane anesthetized female adult Sprague-Dawley rats (∼200g; Harlan Laboratories, UK) received a midline 150 kdyne contusion injury at spinal level T10 using an Infinite Horizon impactor device, as previously described in (15). Control T10 spinal cord tissue was collected from uninjured animals of the same sex, strain and weight. The study has received approval by King’s College London under licence and all surgical procedures were performed in accordance with the United Kingdom Animals (Surgical Procedures) Act 1996. More information in **Supplemental Methods**.

### Protein extraction from spinal cords and shot-gun proteomics

Protein extraction from either T10 injury epicentre or uninjured control T10 spinal cord segments (5–6 mm, ∼45 mg per explant covering the entire injury epicentre) was performed on PBS-EDTA perfused rats using a protein extraction protocol to ensure sequential isolation of cellular (0.08% SDS) and extracellular (4M guanidine) proteins from spinal cords. Liquid chromatography tandem mass-spectrometry shot-gun proteomics (Q Exactive hybrid quadrupole orbitrap) was performed on both 0.08% SDS and 4M guanidine spinal cord tissue extracts from 3 intact and 3 injured (7 days post-SCI) T10 spinal cord segments as previously published in considerable detail (16). Raw MS/MS data was loaded for protein, peptide and spectra validation and visualization onto Scaffold v4 (17). Scaffold was also used to calculate normalised spectral counts for quantitation (18). Protein extraction and proteomics procedures are described in-depth in **Supplemental Methods**.

### Immunoblotting

Protein extracts were electrophoresed and immunoblotted using standard and routine protocols (5, 16). 16μg of protein per sample were separated on 15-well, 4–12% pre-cast gradient gels (NuPAGE, Invitrogen). Proteins were transferred on nitrocellulose membranes, blocked and incubated with 1ry antibodies (1:1000) for 16h at 4^°^C. Membranes were washed and incubated with appropriate horseradish peroxidase-conjugated 2ry antibodies (1:2000; Dako). Finally, enhanced chemiluminescence (ECL) signals were visualised in a UVP BioSpectrum. More information and antibodies used in **Supplemental Methods**.

### Quantitative PCR

Total RNA was extracted from intact and injured T10 spinal cord tissue explants homogenised in Trizol reagent (Sigma) and using routine protocols (5, 16). RNA was purified using the EZNA total RNA kit I (Omega Bio-Tek). 2000 ng of RNA per sample were converted into cDNA using an RNA-to-cDNA kit (Applied Biosystems). 20 ng of RNA per reaction were quantified using rat prevalidated TaqMan primer/probe mixes for CD34, PDGFR, COLA1 and COL1A2 (Applied Biosystems). GAPDH was the housekeeper. Real-time PCR was performed using a Roche thermocycler. Fold-change in mRNA expression of target genes was calculated relative to intact T10 explants using the ΔΔCt method. More details in **Supplemental Methods**.

### Immunohistochemistry

T10 SCI epicentre or intact T10 spinal cord segments were collected from 4% PFA perfused-fixed animals (60min total fixation time). 30μm cryosections were probed with goat CD34 (1:100; RnD), rabbit COL1A1/A2 (1:100; Abcam), rabbit PDGFRα antibodies (1:100; Cell Signalling) and rabbit CD45 (1:100; Abcam). Sections were incubated for 16 h at 4°C and received either donkey anti-rabbit AlexaFluor 568 or donkey anti-goat AlexaFluor 488 for 90 min. Sections were mounted using VectaStain and imaged on a Zeiss Axioplan 2 microscope. More details in **Supplemental Methods**.

### Statistical and Computational analysis

Differential protein expression was measured using spectral counting (Scaffold v4). Differential protein levels were estimated by unpaired, two-tailed T-Test for each identified protein. No outliers were removed. Hierarchical clustering and heat-maps were made in TM4 MeV v4.8 (19). Protein-protein interaction networks were created using StringDB v10 (20). Protein networks were visualised in CytoScape v2.8 (21) and analysed using BinGO (22) to identify overrepresented Biological Process GO terms. Densitometry and TaqMan qPCR data were analysed using T-Test. GraphPad Prism v6 was used to carry out all statistical tests and obtain *p* values. Additional details in **Supplemental Methods**.

## RESULTS

### Rat and axolotl transcriptomics microarrays comparison

To identify molecular differences and similarities between rats and axolotls we performed an intersection of differentially regulated genes identified in a rat SCI microarray (14) 7 days after injury (**Supplemental Table 1**) with differentially regulated genes identified in a microarray dataset of axolotl mexicanum SCI (9) also 7 days post-injury (**Supplemental Table 2**). Both datasets are freely accessible via the gene expression omnibus (GEO-NCBI; Rat: goo.gl/Hz4Gj9 and Axolotl: goo.gl/Eoshgn). Microarray data from GEO is qualitatively and statistically curated and MIAME-compliant (23). Significantly upregulated and downregulated genes from rat and axolotl microarrays are also visualised and qualitatively analysed as protein-protein interaction networks in **Supplemental Fig. 1** and **Supplemental Fig. 2** respectively.

**Figure 1:**
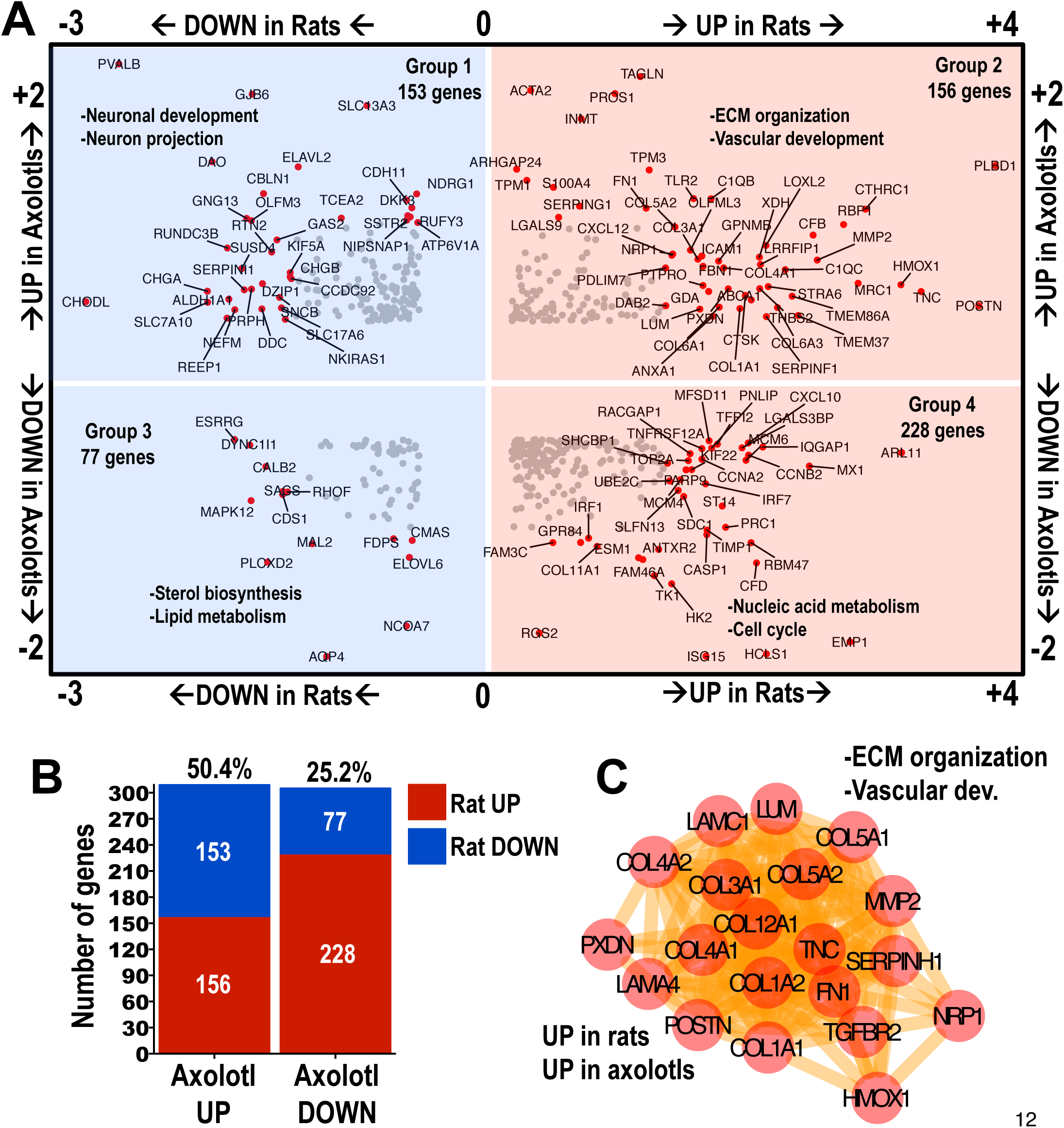
Comparing rat and axolotl spinal cord injury microarrays to identify common differentially regulated genes. **A**: The 4-direction scatter-plot depicts shared differentially regulated genes identified in rat and axolotl microarrays 7 days post-SCI. Only transcripts with adjusted *p* value ≤0.05 and fold-change ≥2 (see **Supplemental Tables 1** & **2**) were included. Shared differentially regulated transcripts were split into 4 groups according to their log fold-change in rat and axolotl microarrays: *Group 1* genes were upregulated in axolotls but downregulated in rats; *Group 2* genes were upregulated in both species; *Group 3* genes were downregulated in both species and *Group 4* genes were upregulated in rats but downregulated in axolotls. Highly differentially regulated genes across species are indicated (red and text) and significant biological process GO terms found using BinGO are shown. **B**: The graph summarises the number of shared genes from intersection of rat and axolotl microarrays 7 days post-SCI and their group distribution. **C**: Protein-Protein interaction network depicting the shared cluster of genes with GO terms “ECM organization” and “vasculature development” from *Group 2*. These genes are upregulated in both rats and axolotls 7 days after SCI. Network was made using StringDB and Cytoscape.

**Figure 2:**
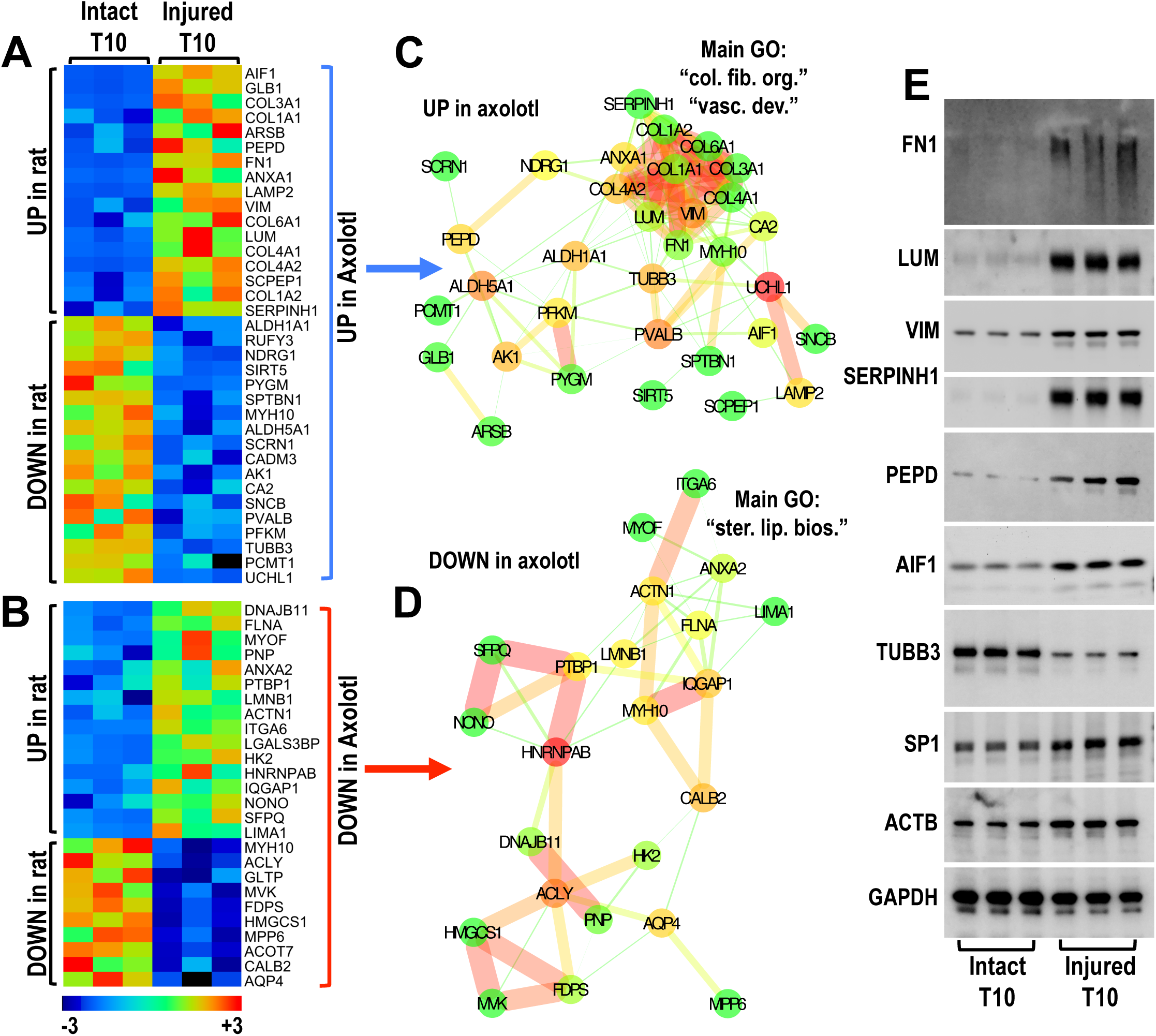
High-throughput proteomics validates the differential regulation of multiple transcripts at the protein level following rat SCI. **A-B**: Heat-maps summarize significantly altered (T-Test *p≥0.05*) proteins identified by shot-gun proteomics and quantified by spectral counting, 7 days after rat SCI (**Supplemental Tables 3** & **4**). These proteins had matching upregulated and downregulated transcripts in the rat microarray and were either upregulated (**A**) or downregulated (**B**) in the axolotl microarray. **C-D**: Protein-protein interaction networks of axolotl upregulated (**C**) or downregulated (**D**) intersecting proteins. **E**: Multiple proteins from **A** & **C** were validated using immunoblotting in intact or injured T10 rat spinal cord extracts 7 days post-SCI.

Next, we compared the datasets to find transcripts that were shared and significantly altered in both species at 7 days post-SCI. Only genes that had adjusted *p* value of ≤0.05 and fold-change ≥2 were included (**Supplemental Tables 1 & 2**). We found 4 groups of shared genes (**Fig. 1A**): *Group 1* with 153 genes upregulated in axolotls but downregulated in rats; *Group 2* with 156 genes that were upregulated in both axolotls and rats; *Group 3* with 77 genes downregulated in both axolotls and rats; *Group 4* with 228 genes downregulated in axolotls but upregulated in rats. While rats and axolotls share 156 upregulated genes (**Fig. 1B**; 50.4% similarity), only 77 genes are downregulated in both species, while 228 of the genes that are downregulated in axolots are instead overexpressed in rats (**Fig. 1B**; 25.2% similarity). We then used BinGO (22) to identify overrepresented biological process gene ontology (GO) terms in each group of shared genes. Genes present in each group together with overrepresented GO terms are presented in **Supplemental Fig. 3**. *Group 1* was characterised by “neuronal development” and “neuron projection” terms (**Fig. 1A** & **Supplemental Fig. 3A**) perhaps reflecting that multiple neuronal transcripts are downregulated in rats but instead are increased in regenerating axolotls after SCI. *Group 2* (upregulated in both species) was enriched in “extracellular matrix organization” (**Fig. 1A** & **Supplemental Fig. 3B**) indicating that matrix remodelling is involved in wound healing in both species. This complex group contained multiple collagen genes (including collagen-1; COL1A1 & COL1A2), but also “response to wounding” (mainly inflammatory genes), “cell adhesion/motility” and “vascular development” terms (**Supplemental Fig. 3B**). *Group 3* (genes reduced in both species) was characterised by “sterol/lipid metabolism” terms (**Fig. 1A** & **Supplemental Fig. 3C**). Cholesterol synthesis is reduced during peripheral nerve regeneration (24). *Group 4* was enriched in “nucleic acid metabolism” and “cell cycle” terms (**Fig. 1A** & **Supplemental Fig. 3D**), perhaps unsurprisingly given the importance of cell cycle regulation in axolotls regeneration (25).

**Figure 3:**
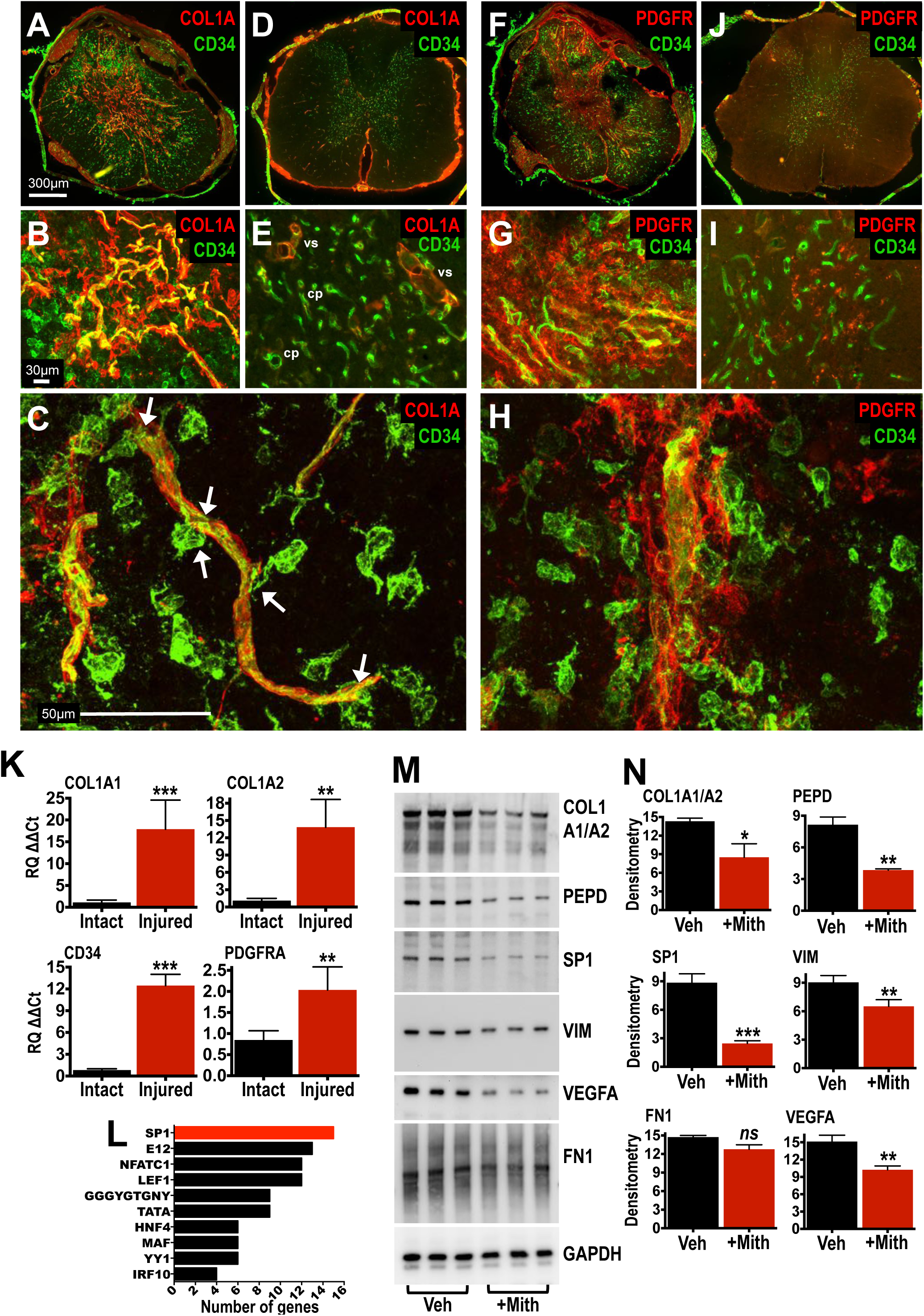
Collagen-1 signature and aberrant vascular remodelling after rat SCI. **A-E**: Immunofluorescence staining of collagen-1 (red; antibody recognises both COL1A1 and COL1A2 chains) and CD34 (green) in either T10 injury epicentre sections (**A**) 2 weeks post-SCI or intact control T10 spinal cord (**D**). (**B)** is a magnified area from (**A**) and (**C**) is an image taken with a 60x oil objective. (**E**) is a magnified area from (**D**). **F-I**: Immunofluorescence staining of PDGFRα (red) and CD34 (green) in either T10 injury epicentre sections (**F**) or intact T10 spinal cord (**J**). (**G**) is a magnified area from (**F**) and (**H**) is a 60x oil objective image from (**F**). (**I**) is a magnified area from (**J**). Results are representative of at least 4 different immunostained spinal cords. **K**: TagMan qPCR of intact and injured spinal cord segments 2 weeks post-SCI. ** p≥0.01, *** p≥0.001 T-Test; 6 intact vs 6 injured spinal cords. Data is mean +SD. **L**: 10 Transcription factors with most predicted DNA binding sites for genes shown in **Fig. 2A-B**. Number of likely target genes is shown in the *x axis*. **M-N**: Immunoblotting and densitometry of selected proteins potentially regulated by SP1 in injured T10 spinal cord tissue extracts. Rats were IP injected daily with either vehicle solution (Veh; 10% DMSO in saline) or 0.2mg/kg mithramycin A (+Mith). Animals were sacrificed 2 weeks post-SCI. * p≥0.05, ** p≥0.01, *** p≥0.001 T-Test; 3 vehicle treated vs 3 mithramycin A treated SCI rats. Data is mean +SD

The upregulation of extracellular matrix entities in both rats and axolotls after SCI (*Group 2*), in particular collagen-1 (COL1A1 and COL1A2), other collagens (COL3A1, COL4A1/A2, COL5A1/A2 and COL12A1), matrix and collagen-associated molecules (LUM, FN1, TNC, PXDN, POSTN, SERPINH1 and MMP2; genes highlighted in **Fig. 1C**), was interesting given that matrix accumulation is characteristic of fibrosis, the cardinal feature of tissue pathology and degeneration in most mammalian organs (26). Importantly, the central nervous system is generally considered to be devoid of classic fibrosis and collagen-1, especially in injuries that do not penetrate the meninges (27, 28).

### Proteomics validates the upregulation of the collagen-1 signature in rats

To identify molecules that showed consistent differential regulation after mammalian SCI at the mRNA and protein level, we performed spinal cord proteomics in rats comparing intact T10 spinal cord segments with T10 injury epicentres 7 days post-SCI, using shot-gun proteomics and label-free protein quantitation with spectral counting (18, 29). To achieve high-sensitivity, we used tissue processing previously developed in our lab which allows the sequential extraction and analysis of mainly cellular proteins (detergent-soluble) in 0.08% SDS, followed by the extraction of highly cross-linked extracellular matrix proteins (detergent-insoluble) in 4M guanidine (16, 29, 30). All protein identifications and spectral counts from SDS and guanidine extracts are shown in **Supplemental Table 3** and **Supplemental Table 4** respectively. All raw LC-MS/MS and processed data is publically available in *Mendeley Data for Journals* (SDS: **goo.gl/3xjtjq** & **goo.gl/s8L6La**; Guanidine: **goo.gl/6zE2fP** & **goo.gl/8ZtEUH**). We then compared the proteomics data with the rat microarray. The SDS proteomics dataset (**Supplemental Table 3**) was used for the intersection of all identified tissue proteins excluding extracellular proteins, which were retrieved from the guanidine dataset (extracellular proteins are indicated in **Supplemental Table 4**). Comparison of rat transcriptomics and proteomics returned 110 molecules that were upregulated and 90 molecules that were downregulated both at the mRNA and protein level 7 days after rat SCI, summarised in **Supplemental Fig. 4**.

When these signatures were compared to the axolotl transcriptomics dataset we found 35 common molecules upregulated in axolotls (**Fig. 2A, C**) and 26 common molecules downregulated in axolotls (**Fig. 2B, D**). Heat-maps (**Fig. 2A-B**) display differential expression from proteomics of rat SCI. Again, shared upregulated molecules included multiple collagens and collagen-associated proteins (**Fig. 2C**), confirming the transcriptomics findings (see *Group 2* **Fig. 1A, C**). Interestingly, many of these matrix proteins were recently found by our group using proteomics in advanced (8 week) T10 rat lesions (16). This consistent signature suggests a conserved response in both species that is related to collagen and potentially vascular remodelling and might be associated with regeneration in the axolotl but tissue scarring in the rat. The cluster of common molecules downregulated in the axolotl was associated with “sterol/lipid biosynthesis” (**Fig. 2D**), again confirming the transcriptomics comparison (see *Group 3;* **Fig. 1A**).

Multiple upregulated proteins from the signature were then validated by western-blotting in lesions from a separate cohort of rats 7 days post-injury (**Fig. 2E**). These included classic SCI proteins fibronectin (FN1), IBA1 (macrophage/microglia marker) and vimentin (VIM; astrocytes, fibroblasts and endothelial cells). Other less known proteins included the collagen-associated proteoglycan lumican (LUM) as well as SERPINH1 (Hsp47) and peptidase D (PEPD), both involved in collagen-1 metabolism. Beta 3 Tubulin (TUBB3), a typical neuronal marker, was clearly downregulated after rat SCI as expected (31), but was instead upregulated in axolotls 7 days after SCI (**Fig. 2A**), indicating the difference in neuronal regeneration between the two species.

### Accumulation of collagen-1 in rat lesions

Collagen-1 was examined further in a separate cohort of SCI rats. Immunohistological localization in spinal lesions found a clear upregulation and accumulation of COL1A (antibody recognises both COL1A1 and COL1A2 chains) after SCI (**Fig. 3A-B**) in comparison to intact T10 sections (**Fig. 3D-E**). Collagen was apparent as a dense meshwork in the centre of the lesion core and extending outwards around the central canal. COL1A-stained structures colocalized with CD34 (**Fig. 3B**), a surface marker highly abundant in endothelial cells and endothelial progenitors, enriched in capillaries, microvessels and small proliferating vessels during angiogenesis (32-37). Costained areas showed a disorganised microvessel appearance indicative of abberant vascular remodelling (**Fig. 3B**). CD34 also stained individual globular cells around collagen-stained areas, presumably lesion-infiltrating endothelial progenitors (**Fig. 3A-B**). High-magnification revealed contact between CD34+ cells with collagen-stained microvessels as well as clear endothelial staining (**Fig. 3C**; white arrows). Intact T10 spinal cord sections (**Fig. 3D-E**) had extensive CD34 staining in grey matter capillaries (**Fig. 3E**; *cp*), while COL1A stained larger vessels in white and grey matter (**Fig. 3E**; *vs*).

In consecutive serial sections (30μm apart) the centre of the lesion core, where most of collagen-1 staining was seen (**Fig. 3A-B**), was also immunopositive for PDGFRα (**Fig. 3F-G**), suggesting that collagen might be related to PGFRα+ lesion stromal cells such as pericytes or ependymal cells, which can contributre to scarring in mouse SCI (38). CD34+ cells were associated with PDGFRα+ regions but they did not apper to colocalize (**Fig. 3F-H**). Upregulation of COL1A1, COL1A2, CD34 and PDGFRα was validated at the mRNA level, in a separate cohort of injured animals (**Fig. 3K**). In advanced cavitated lesions (12 weeks post-SCI), collagen-1 staining was extensive (**Supplemental Fig. 5A-B**). Similar to 2 weeks post-injury (**Fig. 3A-B**), COL1A was centrally distributed, colocalised with CD34 and the staining pattern was consistent with erratic microvessels (**Supplemental Fig. 5B**).

### SP1 regulates collagen-1 in rat lesions

Transcription factor promoter binding site analysis (MSigDB-TRANSFAC; 40,41) found that SP1 was the potential regulator of the highest number of collagen-signature genes (16 genes with likely SP1 promoter binding sites; **Fig. 3L**). SP1 was also increased at the protein level following rat SCI (**Fig. 2E**), and in agreement with rat microarray data (**Supplemental Fig. 1H**). In proof-of-concept experiments, the potential involvement of SP1 in collagen regulation was tested using mithramycin (**Fig. 3M-N**), a drug that binds to DNA sequences with guanosine–cytosine (GC) base specificity and interferes with GC-binding transcription factors, such as SP1 (42). Intraperitoneal injections of mithramycin for 14 days following rat SCI (0.2mg/kg; 43,44) reduced the expression of collagen-signature proteins with predicted SP1 DNA-binding sites including COL1A1 and COL1A2, PEPD, VIM and SP1 itself (**Fig. 3M-N**). Mithramycin injections also reduced the prime angiogenic protein, vascular endothelial growth factor A (VEGFA; **Fig. 3M-N**), also known to be regulated by SP1 (45). Fibronectin (FN1) was only marginally affected. Thus, SP1 might regulate matrix remodelling genes after SCI. Nevertheless, we cannot rule out the possibility that other GC-binding transcription factors might also be affected by mithramycin (notably SP3; 46) and may play a role in regulating this gene signature.

## DISCUSSION

Here we performed for the first time a multi-parametric comparison of rat and axolotl spinal cord injury using high-throughput transcriptomics and proteomics. This approach is unique as it directly examined gene expression differences and similarities between a clinically relevant model of mammalian SCI in rats, with a highly regenerative model of SCI in axolotls. The primary comparative analysis as well as the independently validated findings reported here, were based on recently published studies and publically available microarray and proteomics datasets, highlighting the importance of sharing high-throughput data. Amongst multiple interesting findings, the transcriptomics comparison revealed an extracellular matrix collagen-enriched gene cluster that was upregulated in both rats and axolotls after SCI and was subsequently validated using high-throughput proteomics in rats. While expression of collagen-1 has been shown in murine SCI (39), little is known about collagen and fibrosis in clinically relevant rat contusion SCI (28, 47).

In rats, the unexpected accumulation of collagen-1 in the injury site started early and persisted chronically. Rather than loose interstitial fibrosis that typically affects injured mammalian organs, it appears that collagen-1 accumulation was related to aberrant vascular remodelling as collagen-1 colocalised with an abundance of CD34+ cells and disorganised CD34-stained microvessels. CD34+ cells likely represent a population of existing endothelial cells or progenitors that proliferate out of surviving capillaries (CD34 is abundant in grey matter capillaries -34) in the injury epicentre and penumbra, and participate together with collagen-1 to vascular remodelling (32, 48, 49). CD34+ cells might also be infiltrating (50, 51) hematopoietic or myeloid progenitors (52) but this is less likely as we found that they are distinct from abundant CD45+ cells (leukocytes and macrophages/microglia) in spinal lesions (**Supplemental Fig. 6**). Myeloid endothelial progenitors are CD45+ (52).

The origin of collagen-1 in rat contusion SCI lesions is unknown but it might be expressed by perivascular fibroblasts (39), pericytes (53) or scar-forming ependymal cells (38) as previously shown in mice. Phenotypically complex perivascular cells are recently gaining attention as major players of collagen expression and fibrosis in different mammalian organs (54). Importantly, perivascular cells in mice are typically PDGFRβ+ (39, 53). Unfortunately, in our rat SCI model, we were not able to immunohistologically detect PDGFRβ with 3 different antibodies, although PDGFRB mRNA expression was significantly upregulated 2 weeks post-SCI (**Supplemental Fig. 7**). We then tested the other major PDGF receptor (PDGFRα) and found that collagen-1-stained lesion areas were associated with PDGFRα, a marker of perivascular stromal cells (55) and ependymal cells in mice (38) that participate in tissue fibrosis in different systems including the spinal cord (38, 55). PDGFRα is also a classic marker of NG2+ glia. The involvement of these cells cannot be excluded and more work is required to clarify.

In both rats and axolotls collagen and vascular remodelling after SCI might be linked and likely represent a conserved mechanism aimed at wound healing and regeneration, given that blood vessels provide trophic support to injured tissues. Although vascular remodelling is evidently incomplete and fails to support tissue repair in rats, it is presumably highly effective and organised in axolotls and might represent an important factor in their regenerative process (11). Surprisingly, the role and mechanisms of angiogenesis in axolotls have not been studied in detail, even with regards to limb regeneration, and represent interesting areas for future investigations. In mammalian SCI, the role of angiogenesis has been debated, with protective, neutral or even detrimental effects proposed (50, 56-59). Perhaps future repair-promoting strategies should focus not in enhancing angiogenesis but instead in limiting excess collagen-1 deposition (grey matter capillaries are acollagenous) and potentially in organising vessel formation. For instance, organised vascular remodelling was recently shown to be critical for anatomical and functional regeneration in the peripheral nervous system, which regenerates rather effectively. New blood vessels forming in peripheral nerves after injury act as tracks onto which myelinating cells (Schwann cells) are spatially orientated during neuronal regeneration (60).

Another interesting finding was the upregulation of SP1 after rat SCI. The ubiquitous transcription factor was predicted to regulate many genes in the collagen signature and in proof-of-concept experiments, the non-selective SP1 inhibitor mithramycin reduced the expression of collagen-1, VEGFA and others. While these findings are perhaps unsurprising given the role of SP1 in the regulation of thousands of genes including collagen-1, fibrosis and angiogenesis transcripts (45, 61, 62), it is worth noting that its potential function after SCI has not been studied in depth to this date.

In summary, comparison of rat and axolotl SCI transcriptomics revealed that genes involved in matrix remodelling were upregulated in both species. Not only was the excessive accumulation of collagen-1 following rat spinal contusion unexpected but in addition, the potential association of collagen with pathological vascular remodelling is described here for the first time. Collagen accumulation and vascular remodelling are interesting targets for future therapeutic approaches to mammalian SCI. Finally, our publically-available comparative high-throughput analysis identified a multitude of new genes that were similarly or differentially regulated between the two species and could form the basis for future studies in our field an beyond.

## Author contributions

AD designed, performed and analysed all experiments in this study, conceived and coordinated the study and wrote the paper. KB contributed surgical procedures in rats, imaging procedures (Fig3) and writing of the manuscript. JT contributed Fig1A. BR performed LC-MS/MS procedures and wrote the LC-MS/MS methodology. EJB supervised the study and contributed to experimental design and writing of the manuscript. All authors reviewed results and approved the submitted manuscript.

